# JARA: ‘Just Another Red-List Assessment’

**DOI:** 10.1101/672899

**Authors:** Henning Winker, Nathan Pacoureau, Richard B. Sherley

## Abstract

Identifying species at risk of extinction is necessary to prioritise conservation efforts. The International Union for Conservation of Nature’s (IUCN) Red List of Threatened Species is the global standard for quantifying extinction risk, with many species categorised according to population reduction thresholds. We introduce the Bayesian state-space framework ‘JARA’ (Just Another Red-List Assessment). Designed as decision-support tool, JARA allows both process error and uncertainty to be incorporated into IUCN Red List assessments under criterion A. JARA is implemented via an R package that is designed to be easy to use, rapid and widely applicable, so conservation practitioners can apply it to their own count or relative abundance data. JARA outputs display easy to interpret graphics of the posterior probability of the population trend against the corresponding IUCN Red List categories, as well as additional graphics to describe the timeseries and model diagnostics. We illustrate JARA using three real-world examples: (1) relative abundance indices for two elasmobranchs, Yellowspotted Skate *Leucoraja wallacei* and Whitespot Smoothhound *Mustelus palumbes*; (2) a comparison of standardized abundance indices for Atlantic Blue Marlin *Makaira nigricans* and (3) absolute abundance data for Cape Gannets *Morus capensis*. Finally, using a simulation experiment, we demonstrate how JARA provides greater accuracy than two approaches commonly used to assigning a Red List Status under criterion A. Tools like JARA can help further standardise Red List evaluations, increasing objectivity and lowering the risk of misclassification, with substantial benefits for global conservation efforts.

## Introduction

Quantifying trends in population abundance is central to ecological research and to conservation biology in particular, where understanding extinction risk is necessary to prioritise effort in the face of ever-increasing biodiversity loss [1,2]. Although a number classification protocols exist to assess a species’ extinction risk [3], the International Union for the Conservation of Nature’s (IUCN) Red List of Threatened Species is viewed widely as the global standard [1,4].

To list a species in a threatened category (Critically Endangered [CR], Endangered [EN] or Vulnerable [VU]) on the Red List, expert assessors consider the risks associated with both the small-population paradigm and the declining population paradigm [5] under five assessment criteria (A to E). However, quantitative information on species abundance – as a direct measure of the status of a population – is often preferred over less direct measures of extinction risk, like changes in habitat extent or quality [4,6]. Moreover, the IUCN guidelines allow for assignment to a category (ranging from Least Concern [LC] to Extinct) based only on the criterion that produces the highest estimated risk [4]. Therefore, species are often listed only on the basis of a reduction in population size (Criterion A); as of today > 5,000 species are classified as threatened on this basis alone [7]. Criterion A is thus considered to be the most widely used stand-alone criterion for assigning a Red List status to a wide range of animal taxa, including mammals, birds, reptiles, fishes and insects [2,4,8–10].

Although the IUCN Red List provides a set of unified quantitative decision rules for assigning threatened categories based on population decline thresholds, data quality and analytical approaches can differ vastly [4,6]. Estimating the decline between just two points in time remains widely applied as a rapid assessment approach [6,11,12]. Even when model-predicted abundance indices or population trajectories with associated uncertainty estimates exist, it appears common practice to simply extract the two points that span the required IUCN assessment time frame of the longest period between three generation lengths (GL) or 10 years and then calculate the decline as the ratio between these two points. If the observation period is shorter than three generation lengths, a simple extrapolation formula may be applied [11,12]. Fitting linear or log-linear regression models to timeseries of abundance observations is a second widely-applied approach [6,8,10,11]. An advantage of using a regression is that estimates of the significance of the estimated slope, parameter uncertainty, and goodness-of-fit can be obtained, which is not possible with only two points [6]. However, regression implicitly assumes that the “true” population follows a constant deterministic trend, with all deviations from this trend attributable to statistically-independent observation errors [13]. This assumption does not agree well with our understanding of the real world. For example, a population diminished due to a severe drought year or a spike in poaching will start from a lower state than would be expected from an average long-term trend. In reality, animal population trajectories can diverge substantially from this deterministic expectation due to persistent variations in environmental regimes or human impacts [14–16].

In contrast to deterministic regression models, state-space models provide a general framework for analysing dynamical systems that considers process (year-to-year variation) and observation (or reporting) error simultaneously [17,18]. State-space formulations add more biological realism by assuming a Markovian process, such that the population size in the next time step is conditioned on its current state, but independent of past and future states [18]. From a statistical perspective, the observations are independent given the unobservable (latent) states of population. This can help in preventing violation of independence in observation errors, which often manifest themselves as serial residual correlations when fitting simple regressions to timeseries data. Given these desirable properties, it is not surprising that there has been a rapid uptake of state-space approaches for modelling population dynamics [17,19,20], including more recent applications of quantifying population decline [6,21–23]. State-space models have also been specifically applied in simulation studies that highlight the need for a more rigorous quantification of process and observation uncertainties in the context of threat category (mis)classification [3,6,16].

Bayesian implementations of state-space models offer a powerful framework to improve the characterization and communication of uncertainty during IUCN Red List assessments [24]. The posterior probabilities provide an intuitive and transparent way to express uncertainty about population declines to conservation practitioners [22,25], which can be translated directly into probabilistic statements about a population falling into a threatened category [22,23]. However, developing customized Bayesian state-space models can be technically demanding and time consuming, especially when dealing with often case-specific, ‘noisy’ abundance data that are subject to missing values, irregular spacing or multiple indices that are measured at different scales. These issues may therefore dissuade some conservation practitioners and hamper broader applications.

To address this, we have developed the Bayesian state-space framework ‘Just Another Red-List Assessment’ (JARA). Designed as an easy to use, rapid and widely applicable decision-support tool, JARA allows both process error and uncertainty to be incorporated into Red List assessments. A key output of JARA is an easy to interpret graphic in which the probability distribution of the population decline is displayed against the IUCN Red List categories, and where each category is assigned a probability given process and observation uncertainty. To ensure a high degree of transparency, reproducibility and to allow further development, we provide the R package JARA on the global open-source platform GitHub (https://github.com/henning-winker/JARA), so that JARA can be readily applied by conservation practitioners to their own count or relative abundance data. On GitHub, we provide a number of worked examples. Here, we illustrate the main features of JARA using three real-world examples and a simulation experiment to examine how accurately JARA classifies complex population declines under criterion A, relative to the regression and “two-points” approaches discussed above.

## Methods

JARA is a generalized Bayesian state-space decision-support tool for trend analysis of abundance indices with direct applications to IUCN Red List assessments. The name ‘Just Another Red-List Assessment’ acknowledges JAGS (‘Just Another Gibbs Sampler’, [26]), the software used to run the Bayesian state-space model application. The name reference, together with user-friendly R interface and modulated coding structure follows the example of the new open source fisheries stock assessment software ‘Just Another Bayesian Biomass Assessment’ (JABBA, [27]). JARA enables analysis of one or multiple abundance indices simultaneously, where each index can have a different scale, contain missing years and span different periods. JARA provides the option for fitting relative abundance indices to estimate a mean trend or absolute abundance indices (from e.g. different subpopulations) to produce a summed population trend for the total population.

### State-Space Model Formulation of JARA

A central assumption of the state-space approach is that the abundance (*I*_*t*_) trend follows a Markovian process, such that *I*_*t*_ in year *t* will be conditioned on *I*_*t−1*_ in the previous year. For generality, it is assumed that the underlying population trend follows a conventional exponential growth model *I*_*t+1*_ = *I*_*t*_*λ*_*t*_, where *λ*_*t*_ is the growth rate in year *t* [18]. The growth rate *λ*_*t*_ can vary annually to accommodate fluctuations in reproductive success and survival as a result of environmental conditions, anthropogenic activity or other latent (unobservable) impacts. State-space models are hierarchical models that explicitly decompose an observed time-series into a process variation and an observation error component [21]. On the log scale, the process equation becomes *μ*_*t+1*_ = *μ*_*t*_ + *r*_*t*_, where *μ*_*t*_ = log (*I*_*t*_) and *r*_*t*_ = log (*λ*_*t*_) is the year-to-year rate of change, with variations in log-growth rates following a random normal walk: 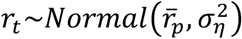 given the estimable process error variance 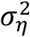 and the estimable mean population rate of change 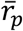 (i.e. the underlying trend). Because the process error is lognormally distributed, *r*_*t*_ is adjusted for lognormal bias by subtracting half the variance (otherwise the stochastic lognormal error induces a small positive bias) at each time step: 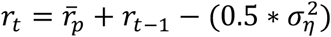 [28]. The corresponding observation equation is of the form log (*y*_*t*_) = *μ*_*t*_ + *ϵ*_*t*_, where *y*_*t*_ denotes the abundance observation for year *t*, *ϵ*_*t*_ is observation residual for year *t*, which is assumed to be normally distributed on log-scale 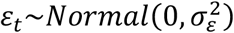 as a function of the observation variance 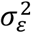.

The model notation for relative abundance indices builds on the approach in JABBA for averaging relative abundance indices [27] and assumes that the mean underlying abundance trend is an unobservable state variable. The corresponding observation equation is then modified, such that log (*y*_*t,i*_) = *μ*_*t*_ + log (*q*_*i*_) + *ϵ*_*t,i*_, where *y*_*t,i*_ is the relative abundance value for year *t* and index *i, μ*_*t*_ is the natural logarithm of the mean abundance trend, *ε*_*t,i*_ is the lognormal observation error term for index *i* and year *t*, and *q*_*i*_ is a scaling parameter for index *i*. The abundance index with the chronologically oldest record is taken as a reference index by fixing *q*_1_ = 1 and the other indices are scaled to this reference index, respectively, with *q*_2,…,*n*_ being estimable model parameters. The estimated posterior of the population trend for year *t* is then given by *I*_*p,t*_ = exp (*μ*_*t*_).

To estimate a total abundance trajectory for the ‘global’ population from multiple timeseries of absolute abundance observations, we assume that each absolute abundance index represents a ‘subpopulation’ that may increase or decline independently from other subpopulations. The process equation is therefore modified to *μ*_*t,i*_ = *μ*_*t,i*_ + *r*_*t,i*_, where *r*_*t,i*_ = log (*λ*_*t,i*_) is the year-to-year rate of change specific to index *i* that is assumed to vary around 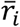 to represent the underlying mean rate of change for the subpopulation (instead of a global population mean 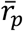), but with a process variance 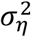 that is common to all subpopulations: 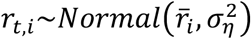. Again, we adjusted *r*_*t,i*_ for lognormal bias by subtracting half the process variance at each time step: 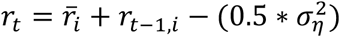 [28]. The corresponding observation equation is adjusted to log (*y*_*t,i*_) = *μ*_*t,i*_+*ϵ*_*t,i*_, so that abundance trend *μ*_*t,i*_ and the error terms 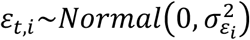 now become the specific to subpopulation *i*. The estimated posterior of the global population trajectory 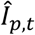 for year *t* is then computed from the sum of all individual subpopulation trajectory posteriors, after correcting them for the lognormal bias by subtracting half their variance [28,29]: 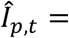 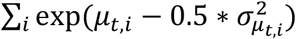. Although the sum of multiple lognormal distributions are not themselves lognormally distributed [30], such that the posterior median or uncorrected mean of 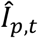 would yield inflated estimates (particularly) when population counts are large [29], our corrected mean is unbiased [31].

### Bayesian Framework

JARA is run from the statistical environment R [32] and executed in JAGS [26], using a wrapper function from the R library ‘r2jags’ [33]. The Bayesian posterior distributions are estimated by means of Markov Chain Monte Carlo (MCMC) simulation. In JAGS, all estimable hyper-parameters are assigned to a prior distribution. JARA uses vague (uninformative) prior distributions throughout, so all inferences are drawn from the information in the data. The estimation of annual growth rate deviates *r*_*t*_, is implemented through hierarchical priors [34], where *r*_*t*_ is informed by the population mean 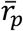 for relative abundance indices and *r*_*t,i*_ is informed by 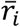 for absolute abundance indices. Vague normal priors with a mean of 0 and variance of 1000, *Normal*(0,1000), are assumed for both 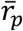 or 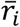. The initial population size in the first year *I*_*t*=1,*i*_ is drawn in log-space from a normal distribution with the mean equal to the log of the first available count *y*_*t*=1,*i*_ and a standard deviation of 1000. Priors for the process variance 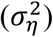 can be either fixed or estimated. If estimated (default), the process variance prior is implemented via a vague inverse-gamma distribution by setting both scaling parameters to 0.001: σ^2^~1/gamma(0.001,0.001) [35–37], which yields an approximately uniform prior on the log scale [27].

The total observation variance 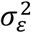 can be separated into three components: (1) externally derived standard errors 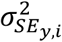 for each abundance index *i*, (2) a fixed input variance 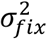 and (3) estimable variance 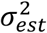, where the prior for 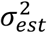 assumes an uninformative inverse-gamma distribution with both scaling parameters set to 0.001. All three variance components are additive in their squared form [38], so the total observation variance for abundance index *i* and year *y* is: 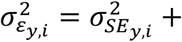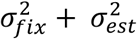 [39]. Each variance component can be individually switched on or off by the user [27]. Adding a fixed observation error is common practice to account for additional sampling error associated with abundance indices [40]. Setting a minimum plausible observation error in this way informs the estimate of the process variance as a portion of total variance is assigned *a priori* to the observation variance [27] and helps to increase model stability and convergence of state-space models [41].

### Estimating probabilities of population decline

A posterior probability for the percentage change (*C%*) associated with each abundance index can be conveniently calculated from the posteriors of 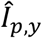, the model predicted population trajectory. If 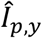 represents a longer time span than the assumed three GL, *C%* is automatically calculated as the difference between the median of three years around the final observed data point *T*, and a three-year median around the year corresponding to *T* − (3 × *GL*). The year *T* + 1 is always projected to obtain a three-year average around *T* to reduce the influence of short-term fluctuations [42]. When the span of 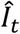 is < 3 × GL, JARA projects forward, by passing the number of desired future years without observations to the state-space model until 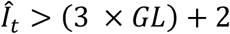. The projections for a single population or multiple subpopulations *i* are based on all possible posterior realizations of *r*_*t,i*_ across all *T* years in the observed timeseries: 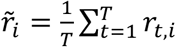

### Model Diagnostics

To evaluate the model fit, JARA provides the user with four plots. The first illustrates the unscaled input data and uncertainty estimates around each observation in the form 95% Confidence Intervals (Figure 1a-b). The second shows the observed and predicted abundance values for each timeseries together with the 95% posterior predictive credibility intervals (Figure 1c-d). The third shows individual fits on the log-scale, as well as the 95% Bayesian credible intervals (CI) derived from the observation variance 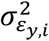 (Figure 2). The fourth is a residual plot (Figure 1e-f) to illustrate potential data conflict when fitting multiple timeseries [27]. This plot includes: (1) colour-coded lognormal residuals of observed versus predicted abundance indices *i*, (2) boxplots indicating the median and quantiles of all residuals available for each year; the area of each box indicates the strength of the discrepancy between the abundance indices (larger box means higher degree of conflicting information), and (3) a loess smoother through all residuals which highlights systematically auto-correlated residual patterns (Figure 1e-f). In addition, the Root-Mean-Squared-Error (RMSE) and the deviance information criterion (DIC) are provided for the comparison of goodness-of-fit and model selection purposes respectively. Convergence of the MCMC chains is diagnosed using the ‘coda’ package [43], adopting minimal thresholds of *p* = 0.05 for Heidelberger and Welch [44] and Geweke’s [45] diagnostics called from R. Unless otherwise specified, JARA reports medians and 95% credible intervals (CI).

**Figure 1:**
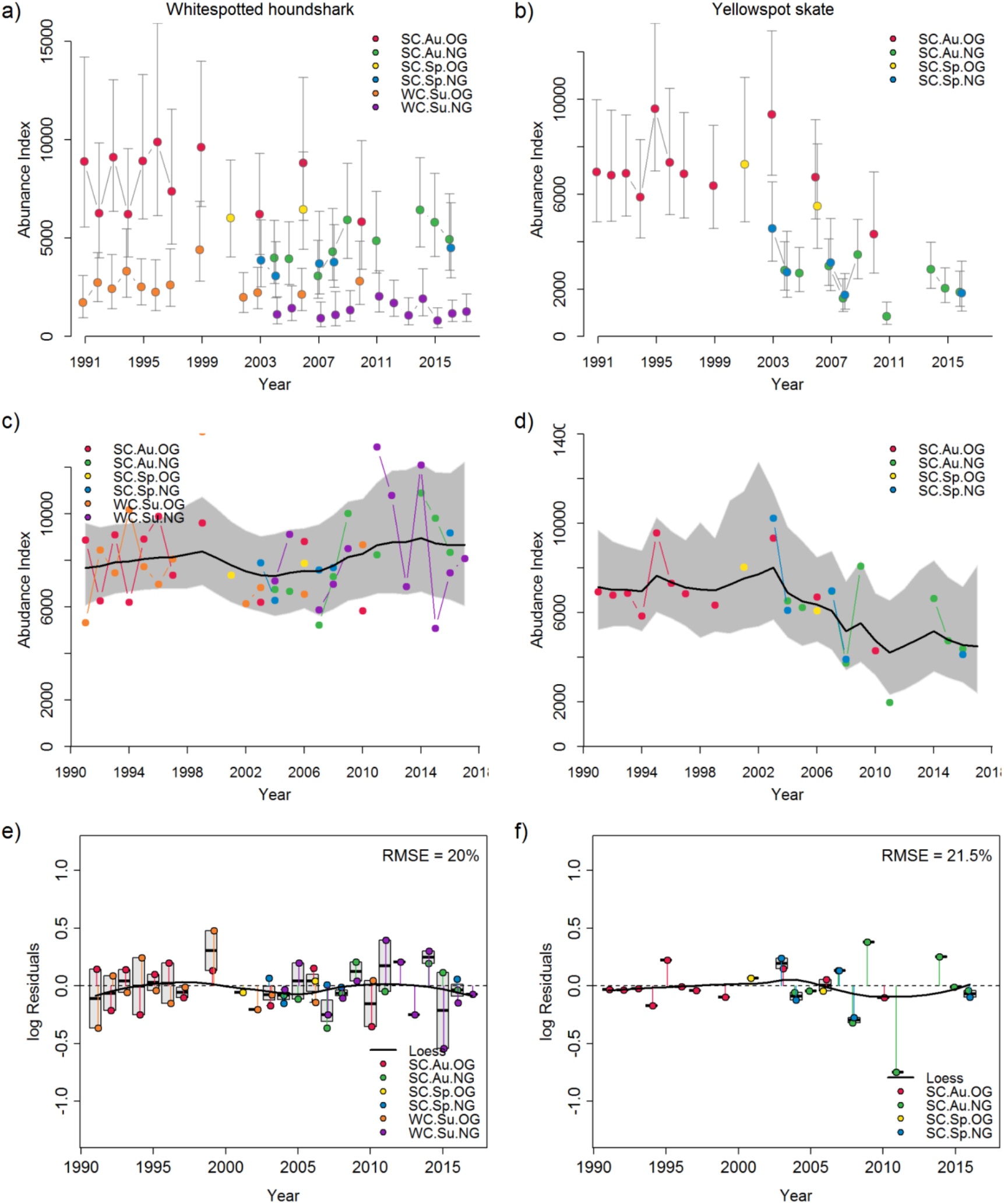
Input datasets of standardized relative abundance indices (± 95% CIs) from South African trawl surveys for two chondrichthyes species, (a) Whitespot Smoothhound *Mustelus palumbes* and (b) Yellowspotted Skate *Leucoraja wallacei* and Model diagnostics plots produced by JARA showing (c) – (d) scaled observed relative abundance (coloured points) for each catch per unit effort (CPUE) timeseries with model fits (predicted relative abundance, black line) and 95% posterior predictive credibility intervals (grey polygons) and (e) – (f) colour-coded lognormal residuals of observed versus predicted abundance indices (coloured points), boxplots (grey bars) indicating the median and quantiles of all residuals available for any given year, and a loess smoother (black line) through all residuals to highlight any systematically auto-correlated residual patterns.

**Figure 2:**
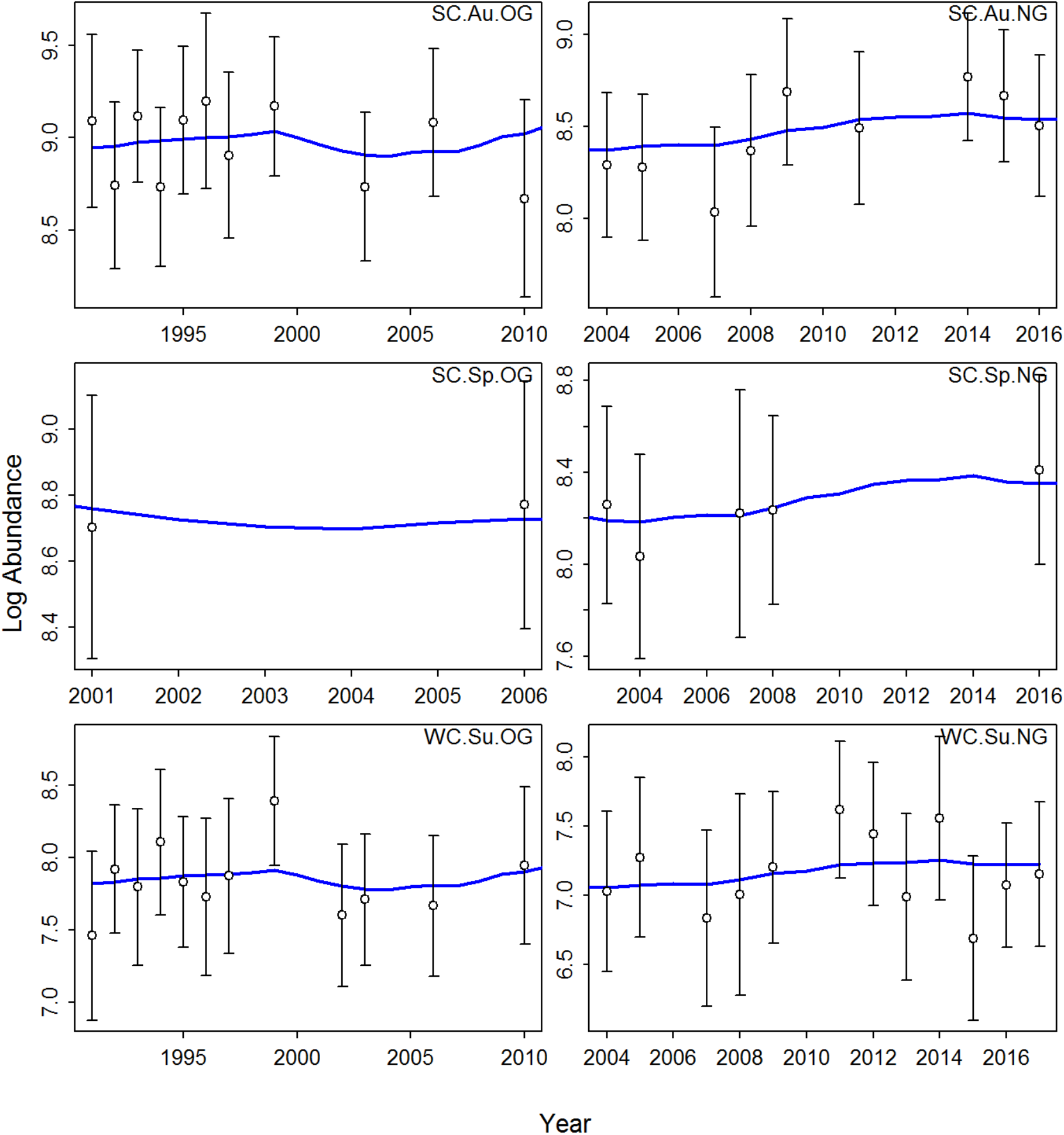
Model diagnostics plots produced by JARA using data from Whitespot Smoothhound *Mustelus palumbes*. Each panel shows the state-space model fit (blue line) to the natural logarithm (log) of the observed relative abundance data (white points), shown with 95% credible intervals (black lines) derived from the observation variance.

### JARA decision support plots

To facilitate decision-making, JARA routinely produces four key plots (Figures 3–5) for each input dataset showing: (1) the combined state-space model fit and 95% CI to abundance timeseries where multiple indices relative abundance (e.g. Figure 4a) and stock assessment results (Figure 4b) are used, or the individual state-space models fits to each count dataset for absolute abundance indices (e.g. individual colony counts for Cape Gannet; Figure 5a); (2) the overall observed and projected (±95% CI) population trajectory over three GL (e.g. Figure 3a and b); (3) the median and posterior probabilities for the percentage annual population change calculated from all the observed data, and from each of the most recent 1 GL, 2 GL, and 3 GL (depending on the length of the observed timeseries), shown relative to a stable population (*%C* = 0) (e.g. Figure 5c); and (4) how the posterior distribution for the percentage change in abundance (*%C*) over 3 GL aligns against the thresholds for the Red List categories (Critically Endangered CR, Endangered EN, Vulnerable VU, Near Threatened NT or Least Concern LC) under criteria A2–A4 (e.g. Figure 4e and f) or under A1. In addition, JARA can be used to undertake a retrospective analysis through the sequential removal of terminal years and subsequent forward projections to attain 3 GL (“retrospective peel”). This enables to user to identify points in time where the *%C* crosses over into different Red List categories (Figure 5f).

**Figure 3:**
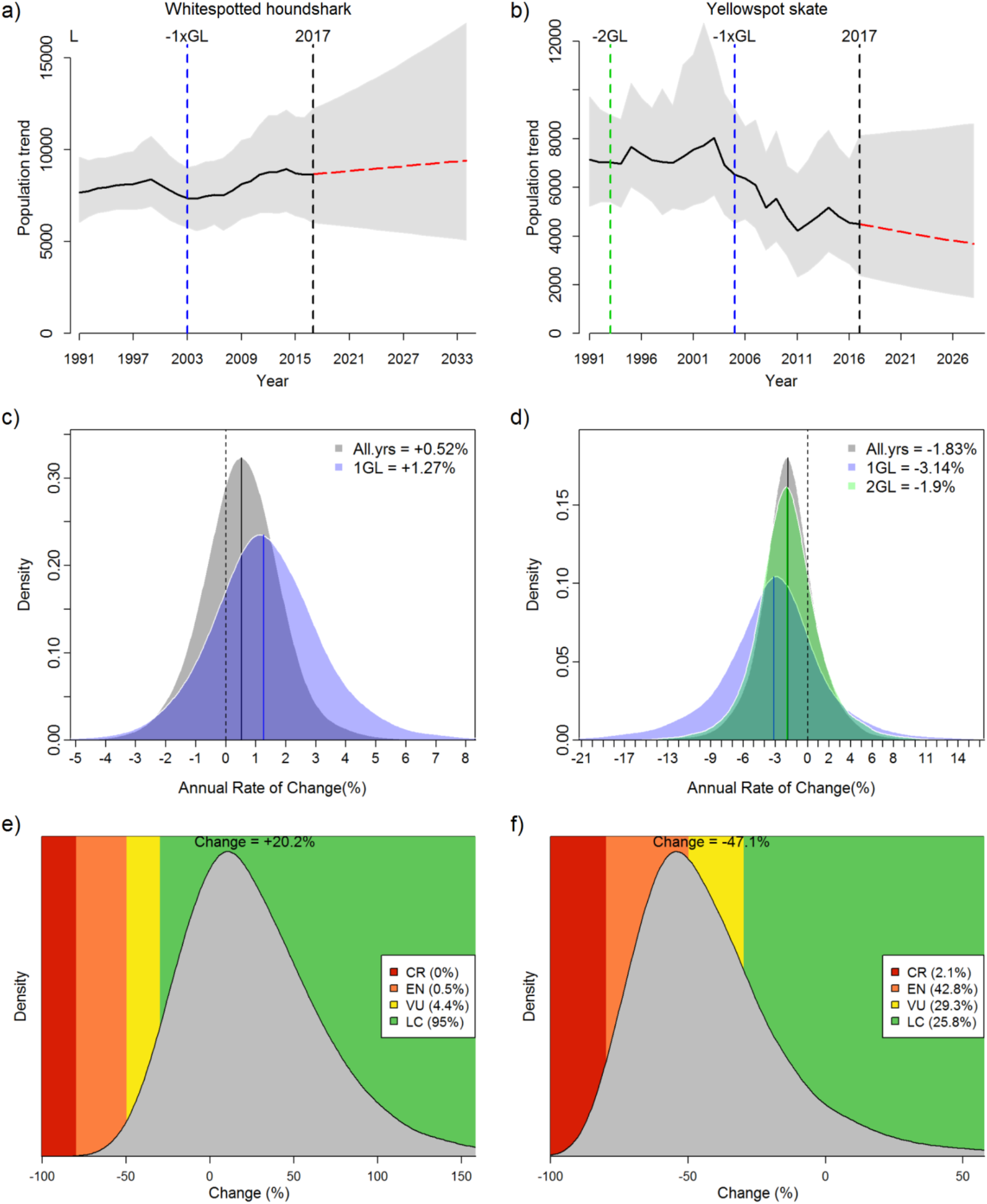
JARA decision-support plots for (left) Whitespot Smoothhound *Mustelus palumbes* and (right) Yellowspotted Skate *Leucoraja wallacei* showing (a) – (b) the overall JARA fit (black line) to the observed timeseries and the projected (red line) population trajectory over three generation length (GL); (c) – (d) the posterior probability for the percentage annual population change calculated from all the observed data (in black), from the last 1 generation length (in blue), from the last 2 generation lengths (in green) with the medians (solid lines) shown relative to a stable population (% change = 0, black dashed line); (e) – (f) the median change over three generation lengths (“Change = xx”) and corresponding probabilities for rates of population change falling within the IUCN Red List categories.

**Figure 4:**
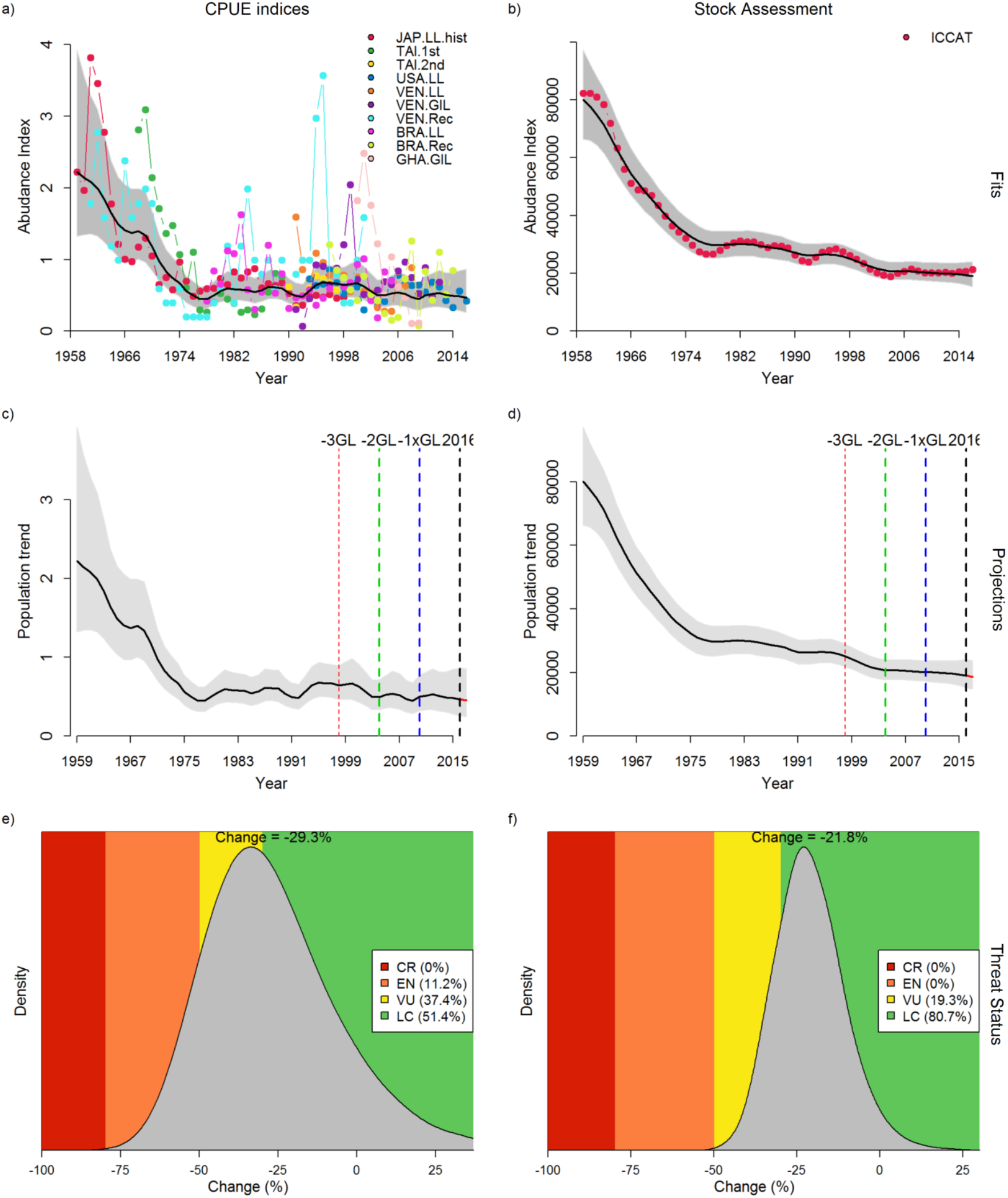
JARA plots for trend analyses of two types of abundance information for Atlantic Blue Marlin *Makaira nigricans*: (left) ten standardized catch per unit effort (CPUE) indices from multiple fishing fleets (1959–2016) and (right) estimated biomass estimates from the 2018 Atlantic stock assessment conducted by the International Commission for the Conservation of Atlantic Tuna (ICCAT), showing (a) – (b) scaled observed relative abundance with model fits (predicted relative abundance, black line) and 95% posterior predictive credibility intervals (grey polygons); (c) – (d) the overall JARA fit (black line) over the observed timeseries with vertical dashed lines denoting the periods corresponding to 1–3 GL; (e) – (f) the median change over three generation lengths (“Change = xx”) and corresponding probabilities for rates of population change falling within the IUCN Red List categories.

**Figure 5:**
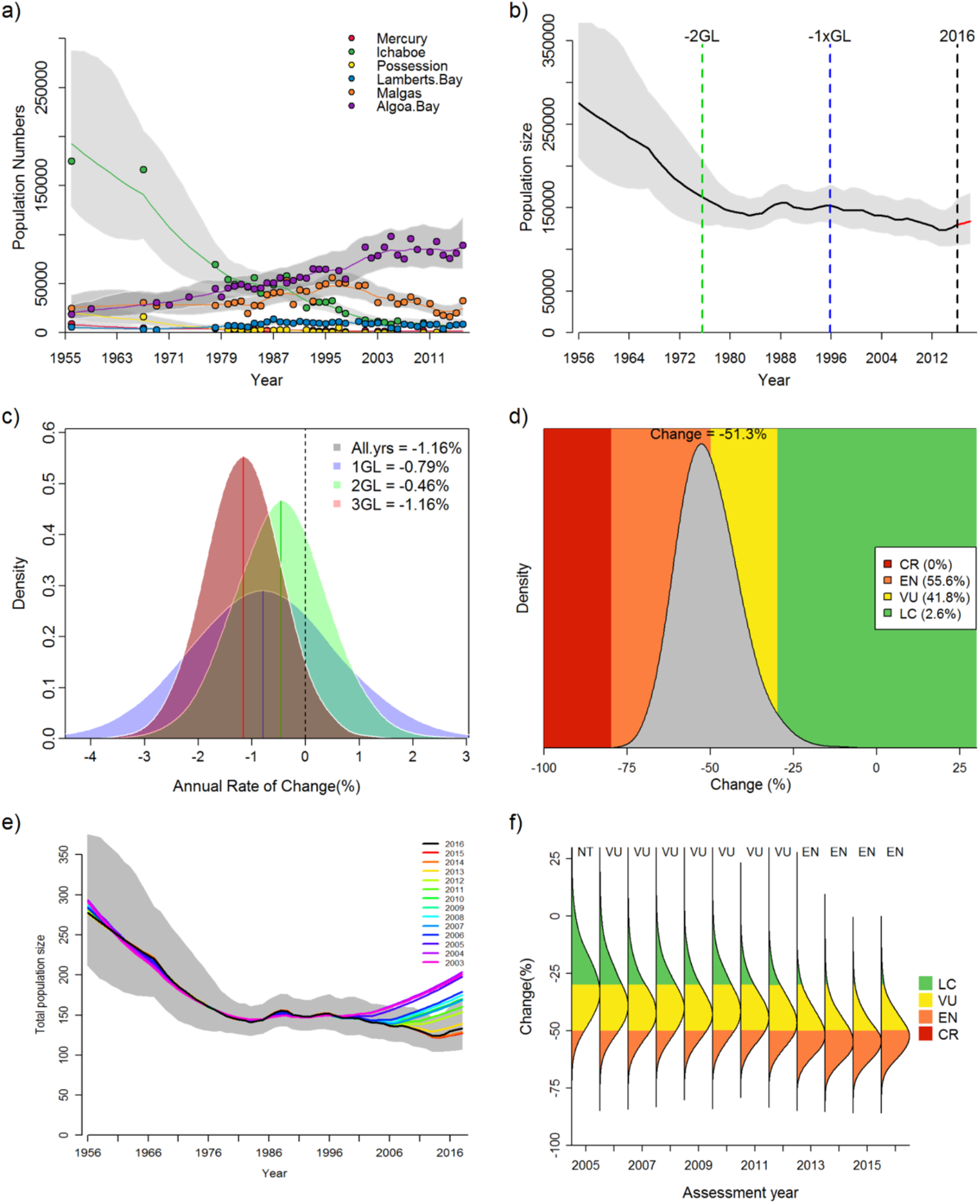
JARA decision-support plots for Cape Gannet *Morus capensis* showing (a) the JARA fits (coloured lines) and 95% credible intervals (grey polygons) to each observed abundance timeseries (coloured points); (b) the overall JARA fit (black line) to the observed timeseries and the projected (red line) population trajectory over three generation times (GLs); (c) the posterior probability for the percentage annual population change calculated from all the observed data (in black), from the last 1 generation length (in blue), from the last 2 generation lengths (in green), and from the last 3 generation lengths (in red) with the means (solid lines) shown relative to a stable population (% change = 0, black dashed line); (e) the median change over three GLs (“Change = xx”) and corresponding probabilities for rates of population change falling within the IUCN Red List categories; (e) retrospective pattern of total abundance estimates obtained through sequential removal of terminal years and subsequent forward projections to attain 3 GL (“retrospective peel”); and corresponding retrospective status posteriors of change over three generation lengths - coloured according to the IUCN Red List category thresholds for the Red List criteria A2.

### Simulation experiment

We conducted a simulation experiment to compare the performance of JARA against two conventional approaches suggested in the IUCN Red List guidelines: (1) the ‘two-point’ approach that calculates %*C* between two years with observations that are 3 × GL apart and (2) regression analysis that extrapolates %C from the estimated slope over a given assessment horizon of 3 × GL [11]. We first used an Operating Model (OM) to generate a ‘true’ population trend and an ‘observed’ abundance index (Figure S1). The observed abundance index is then passed to the three estimation models (EMs: ‘Two-Points’, ‘Regression’, ‘JARA’) to determine the likely threat category (Figure 6).

**Figure 6:**
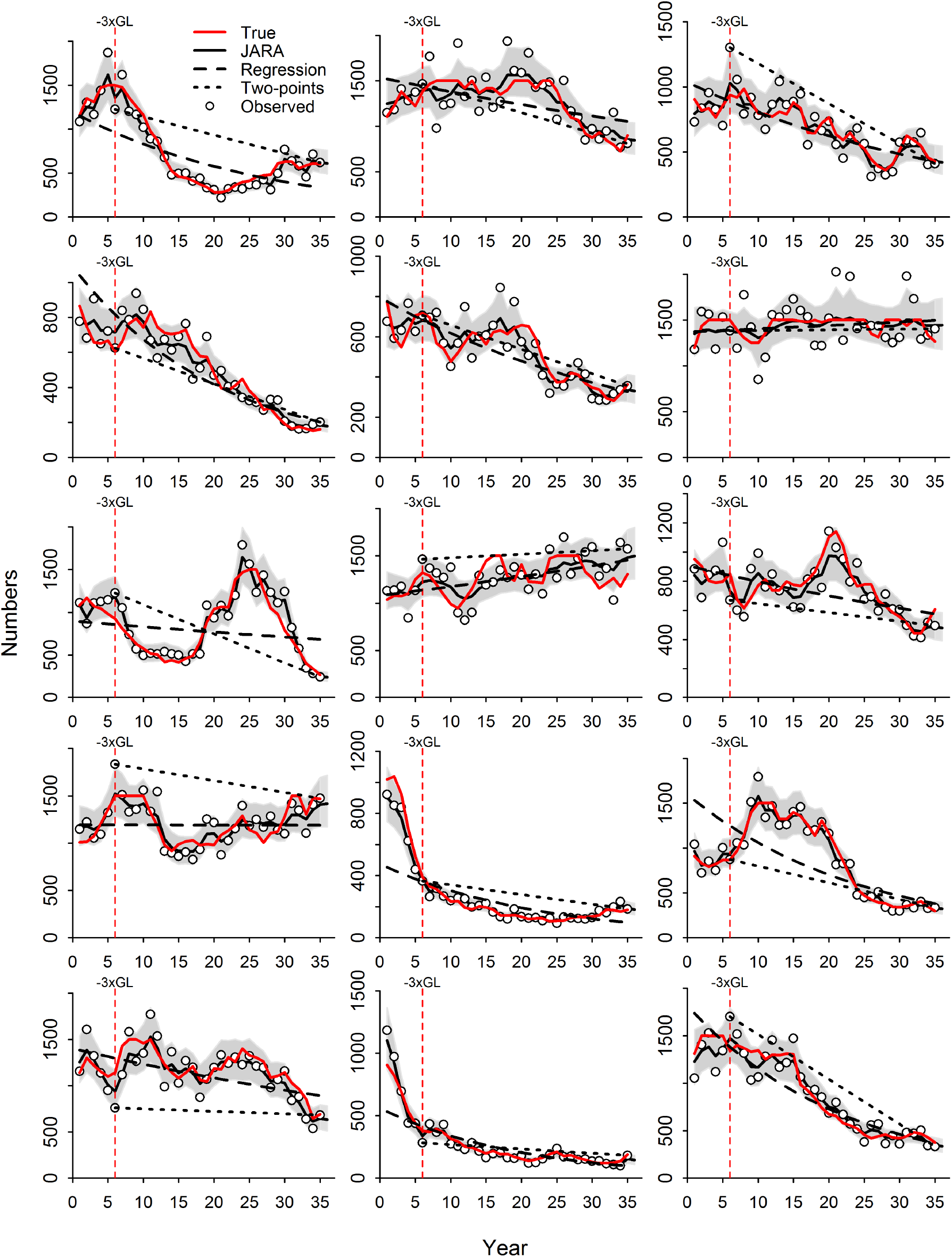
Simulated ‘true’ abundance trajectories and observations for the first 15 of 200 simulation runs together with estimated abundance trends using three alternative estimation models (EMs): (1) ‘Two-Point”, (2) “Regression”, and (3) ‘JARA’. The vertical red dashed line denotes the assessment horizon of 30 years (3 GL) and grey polygons show the 95% credible intervals for the JARA fit.

We modelled the ‘true’ population numbers using a log-linear Markovian process, such that log (*N*_*t*_) = log (*N*_*t−1*_) + *r*_*t*_, where *r*_*t*_ determines the annual rate of change in the population numbers *N*_*t*_. We assumed that *r*_*t*_ can be decomposed into three processes: (i) an underlying mean population trend *r_t_* (ii) stochastic annual variation in the population 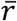 and (iii) annual variation in anthropogenic impact *τ*_*t*_, such that 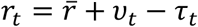 (Fig. S1). The process error deviations associated with *v*_*t*_ and *τ*_*t*_ were assumed to be normally distributed and temporally autocorrelated [15,16] by implementing first-order autoregressive functions (AR1) of the form: 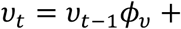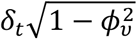 and 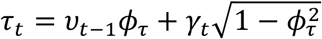 where *ϕv* and *ϕt* are the AR1 autocorrelation coefficients and 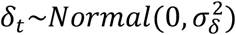 and 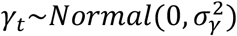 are normally distributed random 9 : variables representing uncorrelated process errors with variances 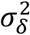 and 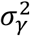, respectively [46]. The 9 : observed abundance index *I*_*y*_ is then generated as a function of the ‘true’ population numbers and a lognormal observation error of the form: 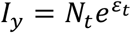, where 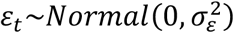 and 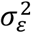 is the an observation variance 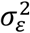. A generic R function of the population simulator is provided in the Supplementary Information.

We simulated 200 random sets of population timeseries of 35 years with an assumed assessment horizon of 30 years corresponding to a GL of 10 years (Figure 6). The initial population size was *N*_1_ = 10,000 and random deviates of 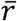, describing the underlying trend, were randomly generated using a uniform distribution between −0.05 and 0 to cover a wide range of population trends and associated *%C* (Figure S1-2). We set the standard deviation for the stochastic variation to a moderate value of *σ*_*v*_ = 0.1 with a process error autocorrelation of *ϕ*_*v*_ = 0.3. This choice of values was intended to describe a population where annual reproductive success is fairly variable and autocorrelation arises from individual age-classes of varying strengths growing through the population. By comparison, the variation in anthropogenic impact was assumed to be less variable *ϕ*_*τ*_ = 0.05 and more persistent over longer, regime-like periods by assuming a fairly large autocorrelation coefficient of *σ*_*v*_ = 0.7. The observation error for the observed abundance was set to *σ*_*ϵ*_ = 0.15 (i.e. CV ~ 15%). The resulting simulations produced a wide range of plausible population trajectories (Figure 6).

We evaluated model performances in terms of the accuracy of the estimated ‘true’ %*C* over 3 GL and the proportion of correctly classified threat categories according to the IUCN Red List A2 decline criterion. Accordingly, the ‘true’ %*C* over 30 years (3 *GL*) was calculated as %*C* = (*N*_35_/*N*_6_ − 1) × 100. For the ‘Two-Point’ EM we calculated 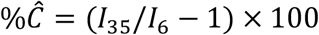. For the ‘Regression’ EM we closely followed the IUCN guidelines by: (i) fitting a log-linear regression of the form log(*I*_*y*_) = *α* + *βy* to all observations *I*_1-35_, (ii) predicting the expected values 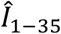, (iii) and then extrapolating the expected 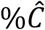 over 3 GL as: 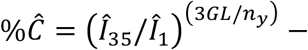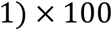 where *n*_*y*_ is the number of years. As described above (*Estimating probabilities of population change*), for the JARA EM the 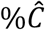 was calculated as the median of the posterior %*C*. Accuracy and bias were calculated using Median Absolute Error (MAE) and Median Error (ME), where the error is the difference between the estimated 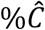 and the ‘true’ %*C*.

## Results and Discussion

### Case study applications

We illustrate applications of JARA using the following worked examples for different abundance data types: (1) multiple relative abundance indices (1991-2017) from South African scientific demersal trawl surveys for two elasmobranchs, Yellowspotted Skate *Leucoraja wallacei* and Whitespot Smoothhound *Mustelus palumbes*; (2) a comparison of model-predicted biomass trajectories from the 2018 Atlantic Blue Marlin *Makaira nigricans* stock assessments and standardized catch-per-unit-effort (CPUE) abundance indices from various fishing fleet, which were used as input into the assessment model and (3) census data for a large seabird, the Cape Gannet *Morus capensis*, in the form of total breeding size estimates for its six global breeding localities in Namibia and South Africa (1979-2017). All worked examples and input data are available on GitHub (https://github.com/henning-winker/JARA).

(1) Whitespot Smoothhound and Yellowspotted Skate are both endemic to southern Africa and had been previously listed as Data Deficient by the IUCN [47,48]. During the 2018 Southern African IUCN Red List workshop, hosted by the IUCN Shark Specialist Group (SSG), both species were reassessed using JARA as a decision-support tool. Based on available life-history information, the GLs were empirically estimated as 14 years for Whitespot smoothhound and 12 years for Yellowspotted Skate. Long-term abundance indices were obtained from analysis of demersal research trawl survey data that were conducted with alternating trawl gear configurations (old/new gear) during summer along South Africa’s west coast (1991-2017) and during autumn and spring along the south coast (1991-2016) by the Department of Agriculture, Forestry and Fisheries. Yellowspotted Skate was only encountered during South Coast Surveys. Annual density estimates (kg per nm^2^ area swept) and associated standard errors on log-scale were estimated using a geostatistical delta-GLMM [49,50]. We only considered the period from 1991 onwards due to evident learning effects for identifying Chondrichthyes during initial survey years. The estimated abundance indices were separately derived for each gear type, survey (coast/season). The input data are characteristic of multiple indices with irregular spacing due to missing survey years (Figure 1 and 2), and are estimated at varying spatial scales. They are therefore treated as relative abundance indices.

JARA combined six abundance indices for Whitespot Smoothhound (Figure 1a) and four abundance indices for Yellowspotted Skate (Figure 1b). Both sets of abundance indices spanned ~2 GL, so JARA automatically projected over ~ 1 GL to provide %*C* estimates of +20.2% with 95% of the posterior falling in LC for Whitespot Smoothhound and −47.1% for Yellowspotted Skate, with EN the best-supported category (Figure 3). The plots of the percentage annual population change (Figure 3c and d) nicely illustrate the contrasting situations for these two species, with Whitespot Smoothhound showing an improving population growth rate in the most recent 1 GL (relative to the whole dataset, Figure 3c), while for Yellowspotted Skate the decline appears to have worsened over the last 1 GL (Figure 3d).

(2) The Blue Marlin, one of the most iconic gamefishes, is a highly migratory billfish species distributed circumglobally in mostly tropical waters. The last IUCN assessment in 2010 classified the species globally as VU [51]. The Atlantic population was assessed by fitting linear regressions to biomass trend estimates from stock assessments undertaken by the International Commission for the Conservation of Atlantic Tunas (ICCAT). GL was assumed to be up to 6 years, corresponding to an IUCN trend assessment horizon of 18 years (3 × GL). Here, we use the 2018 ICCAT stock assessment of Atlantic Blue Marlin to illustrate how JARA can objectively convert such stock assessment outputs into corresponding IUCN Red List decline thresholds and threat categories recommendations. The assessment output used as input for JARA is the median of the estimated biomass for the period 1950-2016 across all accepted candidate models (Figure 4b). Uncertainty was represented by annual standard errors (SEs) of the log-biomass estimates [52]. We then compared the JARA results with a complementary JARA analysis of multiple CPUE indices that served as input data into the 2018 stock assessment. This CPUE dataset comprised standardized CPUE indices (+SEs) from ten different fishing fleets that covered different periods between 1959 and 2016 (Figure 4a).

JARA estimated similar median %*C* for the Atlantic Blue Marlin whether using the multiple CPUE indices (−29.3%) or the stock biomass (−21.8%) as input data. In both cases, the best-supported category was LC, but uncertainty was (unsurprisingly) lower when using the biomass estimates from the stock assessment (80.7% of the posterior in LC) versus the much more variable and partially conflicting CPUE indices (51.4% in LC; Figure 4e and f). Although either input series would likely lead to the same outcome (a classification of LC) if JARA were used a decision-support tool to assess Atlantic Blue Marlin, the analysis of the stock assessment data would result in greater confidence in the assessment’s conclusion.

(3) The Cape Gannet is a large seabird endemic to southern Africa, where it breeds during the austral spring and summer at six islands. Aerial photographs and on the ground measurements of nesting density at all six breeding colonies have been used by the Department of Environmental Affairs (South Africa) and the Ministry of Fisheries and Marine Resources (Namibia) to estimate the total breeding population since the summer of 1956/57 [53]. Counts were sporadic until 1978/79 but undertaken approximately annually thereafter (Figure 5). Cape Gannets generally have high adult survival, strong breeding site fidelity [54] and low reproductive success (≤ 0.82 chicks fledged per pair annually; [55]. Accordingly, their GL is estimated as 20.2 years [56] and we use annual count data from 1956/57 to 2016/2017, which spans 3 GL or the recommended Red List assessment period. Of a possible 366 annual counts between 1956/57 to 2016/2017, 173 were not made for various reasons. Thus, the Cape Gannet dataset is chosen an example of multiple absolute abundance indices (with missing data) that in sum represents the trend of the global population.

The output for Cape Gannet nicely exemplifies the decision-support element of JARA. Based on only the median %*C* (−51.3%), the species meets the criteria for classification as EN (Figure 5d). However, the uncertainty spans LC to EN; assessors may, therefore, also want to consider that while 55.6% of the posterior probability distribution falls into EN, 41.8% also falls into VU (Figure 5d). Moreover, the percentage annual population change plots show that the decline has slowed and then accelerated over the last 2 GL (Figure 5c), while the retrospective plots show that the threshold for EN was first exceeded in 2013, but the situation has not worsened markedly to 2016. Assessors could then make an informed decision on whether to list the Cape Gannet as VU or EN, depending on how risk prone or adverse they wanted to be in their assessment and the presence of other mitigating factors.

### Simulation experiment results

Median Absolute Errors (MAEs) between the estimated and ‘true’ percentage change over 3 generation lengths (GL) using JARA as the EM was 3.63%. The Regression EM (7.87%) and Two-Point EM (10.96%) had MAEs >2 and >3 times larger, respectively, showing a distinct decrease in the accuracy of the estimated %*C* in comparison to the JARA EM (Figure 7). In addition, although all three EMs showed a tendency towards a positive bias (Median Errors > 0), this bias was ~3.7 times larger for the regression EM (0.71%) and ~12 times larger for the Two-Point EM (2.26%) than for JARA (0.19%), which was approximately unbiased. Moreover, the median %*C* from JARA correctly identified the ‘true’ IUCN threat category in 89.5% of cases; > 10% more than the regression EM (which was based on the current IUCN guidelines) and >20% more than the Two-Point EM (Figure 7). Although these might seem like relatively small improvements, misclassification on the IUCN Red List can have real consequences for species conservation [7,57], particularly resource allocation if species move in and out of the threatened categories [58]. There will be many real-world cases where the observed or estimated decline is close to a Red List category threshold (e.g. >50% decline for EN), as was the case for Yellowspotted Skate (median %*C* −47.1%, Figure 3f) and Cape Gannet (median %*C* −51.3%, Figure 5d) in our examples. Not only would we would expect JARA to indicate the ‘true’ IUCN threat category more reliably in these sorts of situations, but the additional information provided by JARA – particularly the plot of the probabilities that the rate of change falls within the IUCN Red List categories (e.g. Figure 5d) - offers assessors insight into the weight of evidence underpinning their assignment.

**Figure 7:**
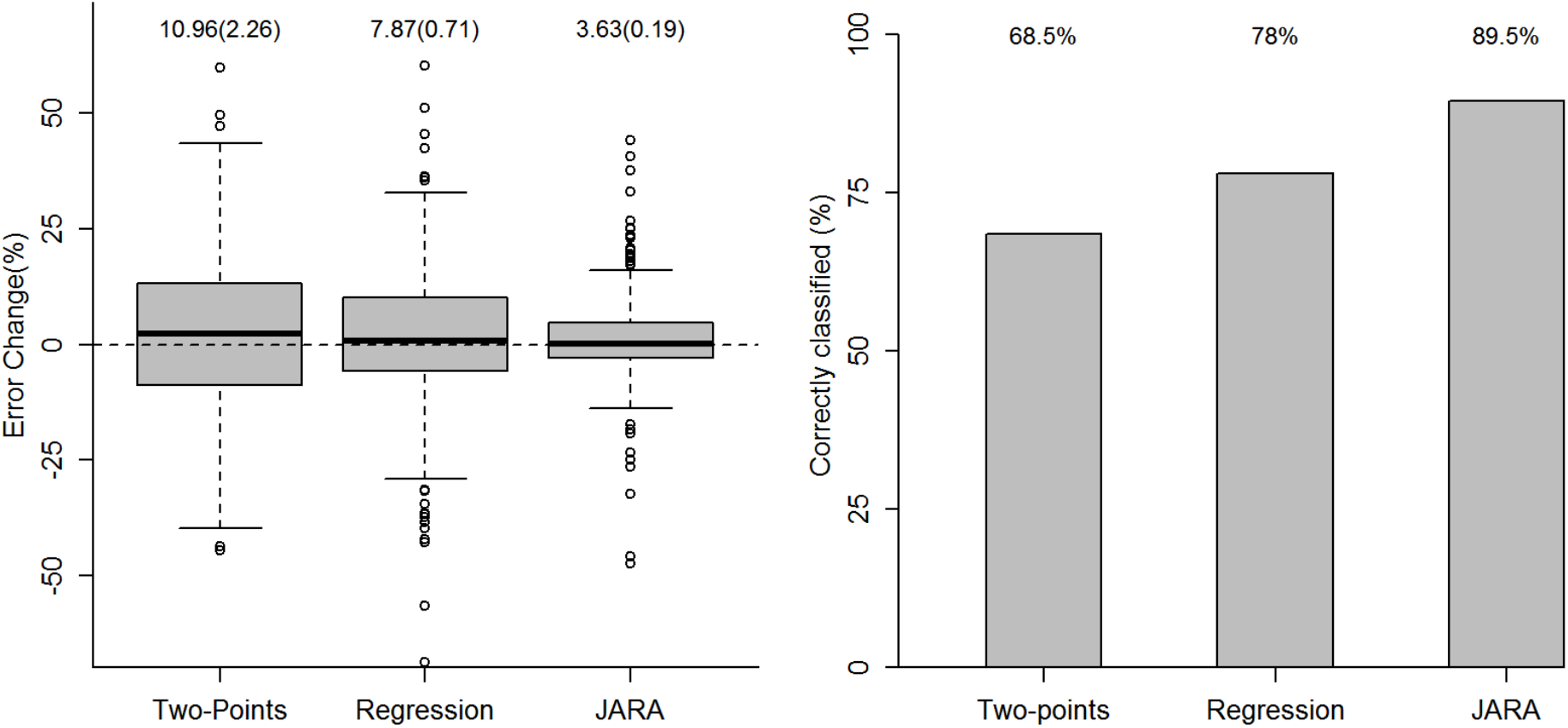
Simulation testing results across 200 runs showing (a) boxplots of errors between the estimated and ‘true’ percentage change over 3 generation lengths (GL) with Median Absolute Errors and Median Errors (in brackets) displayed of each box and (b) barplots illustrating the percentages of correctly classified IUCN threat categories (LC, VU, EN, CR) for the three alternative estimation models (EMs): (1) ‘Two-Point”, (2) “Regression”, and (3) ‘JARA’.

### Current limitations and real-world applications

Although still being refined, JARA has already been used to support expert decision making in the assessment of a terrestrial mammal [59], a number of elasmobranchs [31,60] and two seabirds [31,53]. We designed JARA with the aim of supporting the global effort to provide robust, consistent, and transparent assessments for all species under the IUCN Red List criteria [24]. And we designed it to be free, open-source, and easy to use, to support access for conservation practitioners, researchers and assessors working in places with limited opportunities to access proprietary software [12]. While we certainly hope that more conservation practitioners will use JARA, we caution against applying it to unfamiliar timeseries or blindly accepting the status suggested by decision-support tools [12]. Undertaking Red List assessments requires assessors to understand their target species and the rules specified in the Red List guidelines. The data used for assessments should be proofed and verified by specialists (e.g. an IUCN Specialist Group) to ensure their reliability, users should make informed ecological judgments regarding the length of the timeseries relative to a species’ GL when assessing against criterion A, and should remain cognisant of biases in data availability or survey effort when making their recommendations [11,12,24]. Moreover, JARA has been designed to be used as a decision-support tool and its outputs are not necessarily indicative of a final classification of extinction risk [24]. Rather, assessors should use tools like JARA to make an informed interpretation based on all of the Red List guidelines before coming to a decision on the final assessment outcome [12,24]. Nevertheless, when coupled with expert oversight, JARA has wide potential applicability to other studies of wildlife populations threatened with extinction [61,62].

## Acknowledgments

H.W. and R.B.S. thank Marc Kéry and Michael Schaub for introducing us to Bayesian state-space models, and we thank Nick Dulvy for helpful conversations about the Red List process during the development of JARA. RBS was supported by the Pew Fellows Program in Marine Conservation at The Pew Charitable Trusts. The views expressed are those of the authors and do not necessarily reflect the views of The Pew Charitable Trusts. N.P. was funded by the Shark Conservation Fund, a philanthropic collaborative pooling expertise and resources to meet the threats facing the world’s sharks and rays. The Shark Conservation Fund is a project of Rockefeller Philanthropy Advisors.

## Supplementary Information

**Figure S1:**
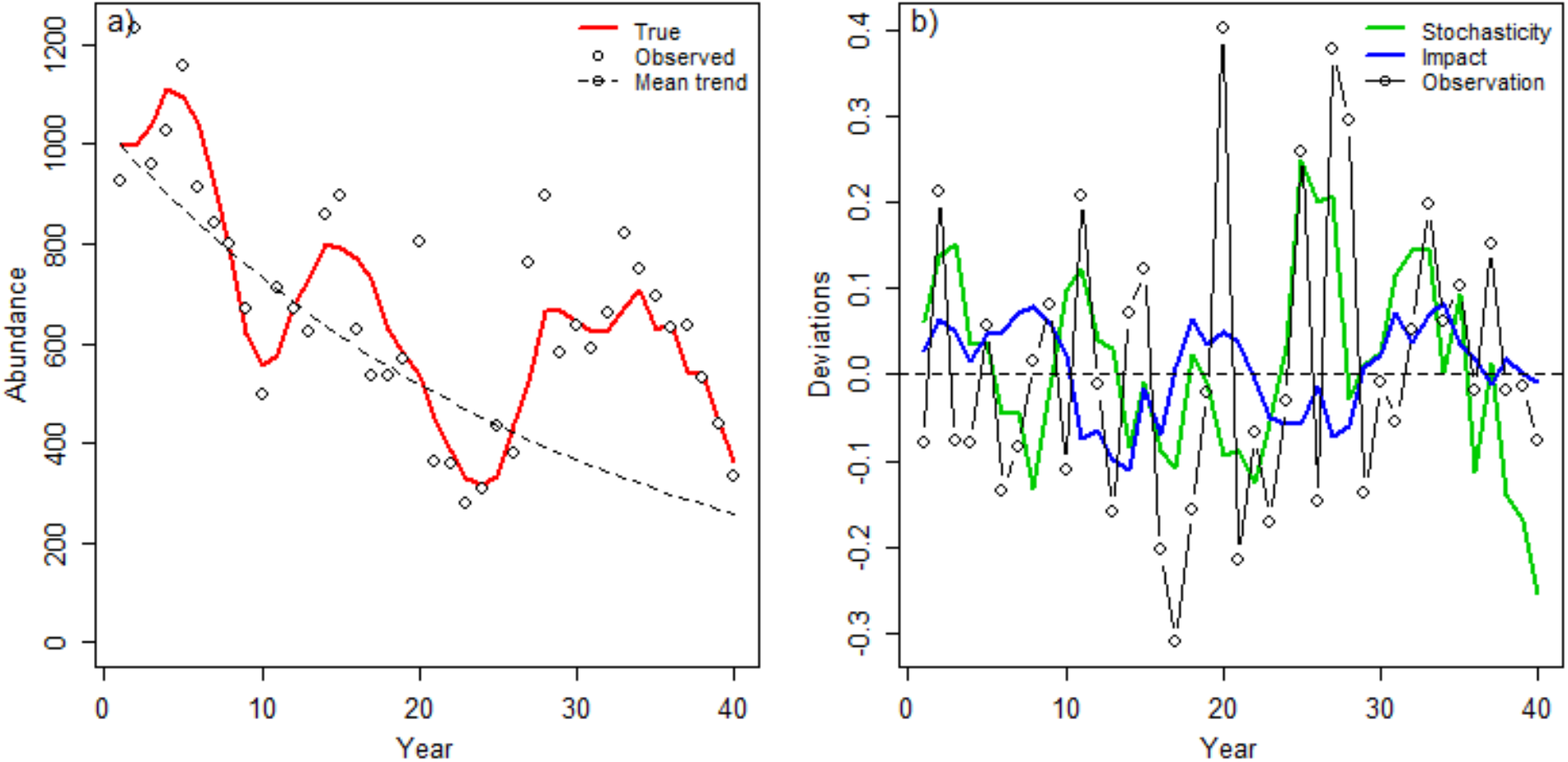
Illustrations of (a) a simulated ‘true’ population trajectory, the corresponding observed values given the observation error and the underlying deterministic mean trend 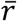; and (b) decomposed first-order autoregressive process error deviations simulating the natural variation in population numbers (Stochasticity) and a more persistent, regime-like variation in anthropogenic impact (Impact), together with a random observation deviations from the ‘true’ abundance due to random sampling error.

**Figure S2:**
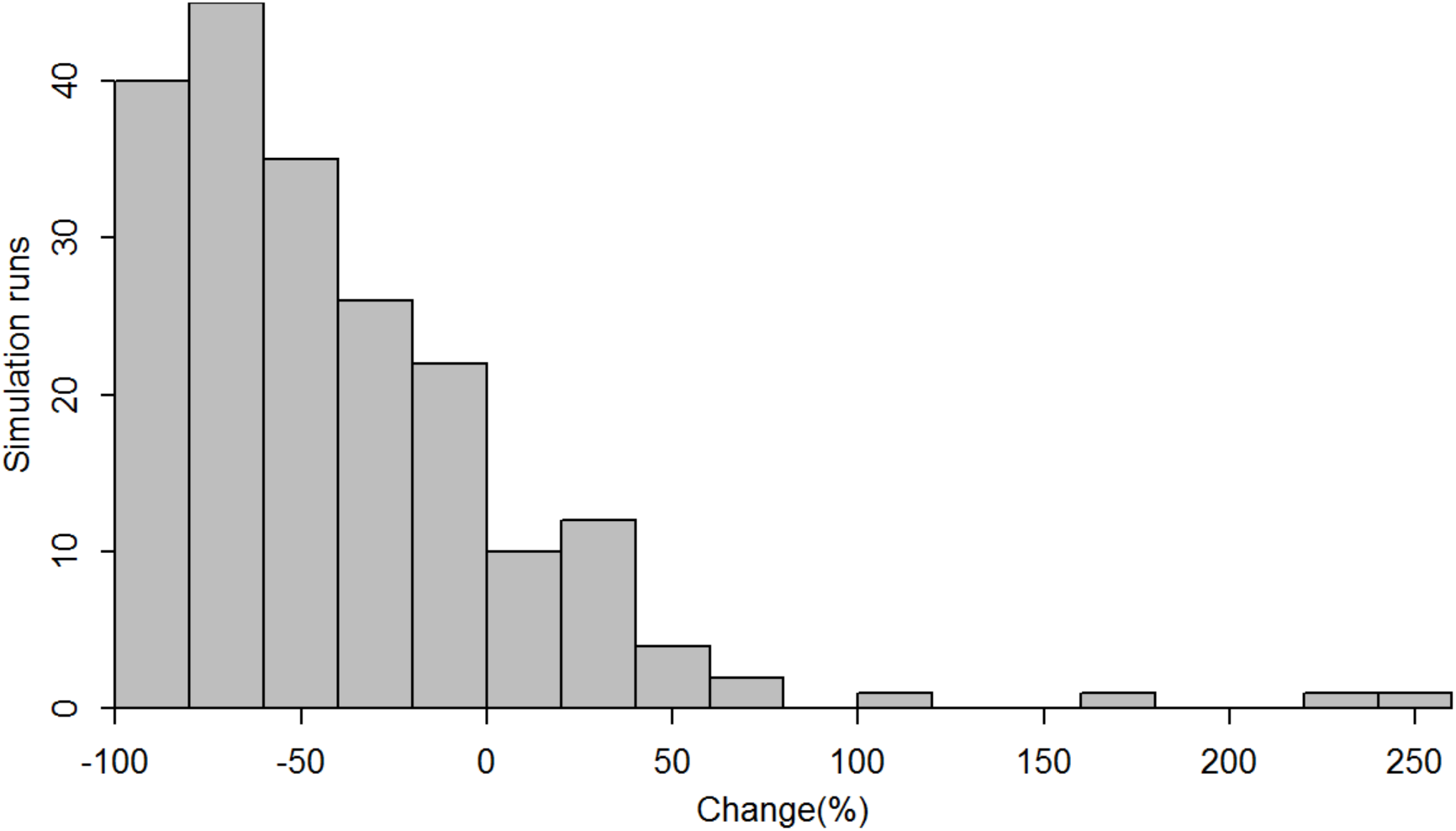
Simulated distribution of the ‘true’ %*C* generated from 200 simulation replicates of abundance trajectories over a 30-year assessment horizon.

---

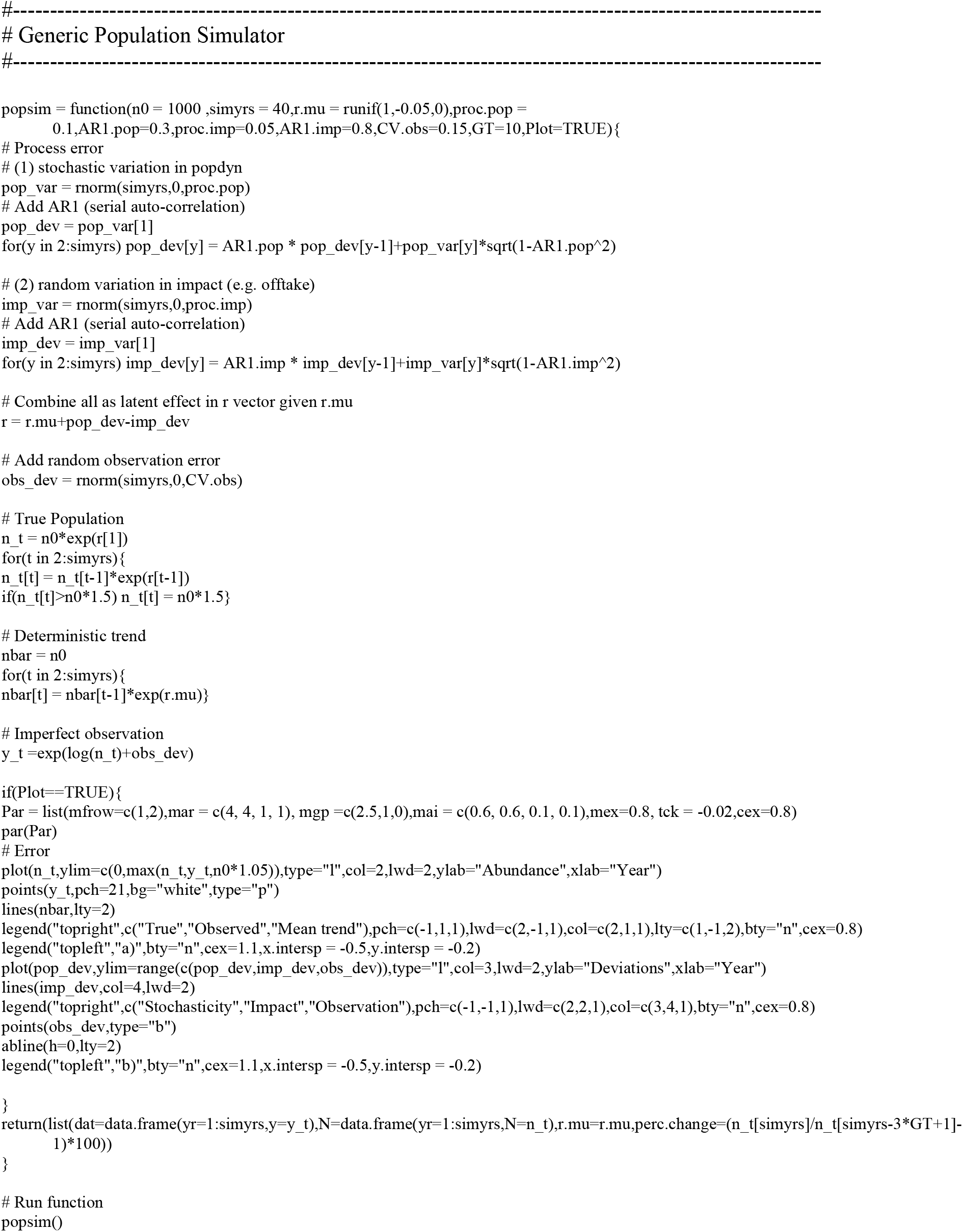

## Notes

https://github.com/henning-winker/JARA

### Competing Interest Statement

The authors have declared no competing interest.

### Summary of Updates

Results and discussion updated, some equations updated, some methods updated, reference made to the JARA R library, new co-author (Nathan Pacoureau) onboard, reference format changed.

